# DenovoProfiling: a webserver for de novo generated molecule library profiling

**DOI:** 10.1101/2021.01.04.425063

**Authors:** Zhihong Liu, Jiewen Du, Bingdong Liu, Zongbin Cui, Jiansong Fang, Liwei Xie

**Affiliations:** State Key Laboratory of Applied Microbiology Southern China, Guangdong Provincial Key Laboratory of Microbial Culture Collection and Application, Guangdong Open Laboratory of Applied Microbiology, Guangdong Institute of Microbiology, Guangzhou 510070, China; Beijing Jingpai Technology Co., Ltd. 1500-1, Hailong Building Z-Park, Beijing, 100090, China; Science and Technology Innovation Center, Guangzhou University of Chinese Medicine, Guangzhou, China; Zhujiang Hospital, Nanfang Medical University, Guangzhou, China; College of Public Health, Xinxiang Medical University, Xinxiang, China

## Abstract

With the advances of deep learning techniques, various architectures for molecular generation have been proposed for de novo drug design. Successful cases from academia and industrial demonstrated that the deep learning based de novo molecular design could efficiently accelerate the drug discovery process. The flourish of the de novo molecular generation methods and applications created great demand for the visualization and functional profiling for the de novo generated molecules. The rising of publicly available chemogenomic databases lays good foundations and create good opportunities for comprehensive profiling of the de novo library. In this paper, we present DenovoProfiling, a web server dedicated for de novo library visualization and functional profiling. Currently, DenovoProfiling contains six modules: (1) identification & visualization, (2) chemical space, (3) scaffold analysis, (4) molecular alignment, (5) target & pathways, and (6) drugs mapping. DenovoProfiling could provide structural identification, chemical space exploration, drugs mapping, and targets & pathways. The comprehensive annotated information could give user a clear picture of their de novo library and could provide guidance in the further selection of candidates for synthesis and biological confirmation. DenovoProfiling is freely available at http://denovoprofiling.xielab.net.

## Introduction

The main objective of drug discovery is to identify a molecule with desired biological properties[1]. Primarily, high throughput screening (HTS) techniques which allow large size of chemical library testing[2,3]. However, HTS is expensive and with low hit rates, only limited to large pharmaceutical companies. Computational based virtual screening methods are used to reduce the size of testing molecules. Various ligand-based[4,5] and structure-based[6] virtual screening methods have been proposed. However, the cost and time consuming for developing a new drug are still increasing[7].

De novo drug design is one of the most promising and scalable approach to address this issue, and in particular, with the advances of deep learning techniques[8–10]. In the early stage, evolutionary algorithms were used for de novo molecular generation[11], which commonly based on the combinations of molecular fragments derived from drug-like library. Over the past years, artificial intelligence algorithms, like deep learning, reinforcement learning, and transfer learning are proposed in the molecule generation field, inspired by the wide applications of those methods to generate text, images, video, and music[12,13].

Recently, a number of architectures for molecular generation, such as recurrent neural networks (RNN)[1,14,15], variational autoencoders (VAE)[16], and generative adversarial networks (GANs)[17] have been developed and proven successful in generating target-focus molecule library. Furthermore, scaffold-constrained molecular generation methods[18,19] were developed for lead optimization. Yang et al. also developed linker constraints molecular generation methods using deep conditional transformer neural networks for fragment-based drug design (FBDD)[20]. Zhavoronkov et al. developed a deep generative model with reinforcement learning and discovered potent discoidin domain receptor 1 inhibitors in 21 days, and further animal experimental results in one lead candidate demonstrated favorable pharmacokinetics[21]. This important case illustrated the utility of deep generative model for the rapid design of compounds with synthetically feasibility, and bioactivity for target of interest. Yang et al. developed a generative model using long short-term memory (LSTM) neural network and generated a focused library containing 672 valid molecules[22]. After filtering with various criteria, synthesis and bioactivity testing, they identified a highly potent inhibitor against p300 with IC50 of 10 nM. These successful cases demonstrate that the deep learning based de novo molecular design could accelerate the drug discovery process.

The flourish of the de novo molecular generation methods and applications created great demand for the visualization and functional profiling for the generated molecules. Generally, the generative models could generate a large chemical library based on sampling criteria and could output with various formats. The following issues, particular for medicinal chemist, are to visualize, analyze, and select the candidates among the generated molecules. Owing to the development of combinatorial chemistry and high-throughput screening technologies, chemical structures and bioactivity data have rapidly accumulated in the past years and are becoming available in public repositories[23,24]. There are various well-established cheminformatics and bioinformatics databases available for drug discovery, which provide comprehensive information for bioactive compounds, drugs, targets, pathways, and disease, like and PDB database[25], PubChem[26], DrugBank[27], ChEMBL[28], and BindingDB[29]. The rising of publicly available databases creates good opportunities for comprehensive profiling of the de novo library.

Dealing with chemical libraries is a common practice in drug discovery. Thus, various cheminformatics tools have been developed for chemical library processing and data analysis. Well-known tools for dealing with chemical library are ChemicalToolbox[30], DataWarrior[31], WebMolCS[32], ChemMine[33], CART[34], MONA[35], and CSgator[36]. Those tools mainly focused on specific functionality, such as large library visualization, structure search or clustering analysis. Even more, some tools are desktop applications, which limited the application. Web based tools dedicated for de novo generated molecule profiling are rare.

In this work, we present the DenovoProfiling, a webserver for de novo generated molecule library profiling. We aim to provide a user-friendly public webserver to support the structure and chemical space visualization, scaffold analysis, molecular alignment, drugs profiling, targets and pathway profiling. We integrated cheminformatics tools and databases to provide comprehensive annotations for the de novo generated molecules. We believe that DenovoProfiling could be an efficient tool for user to capture the knowledge of de novo generated molecules quickly. DenovoProfiling is freely available at http://denovoprofiling.xielab.net.

## Methods

### Framework

The framework of DenovoProfiling was outlined in **Fig. 1**. We integrated the well-known public database PubChem, ChEMBL, DrugBank, and employed open-source cheminformatics toolkit RDKit, and other tools to provide comprehensive information for user submitted de novo chemical library. The profiling process is fully automatically, which user only submit its de novo library files with multiple formats are supported. DenovoProfiling contains 6 modules: identification & visualization, chemical space, scaffold analysis, molecular alignment, target & pathways, and drugs mapping.

**Fig. 1.**
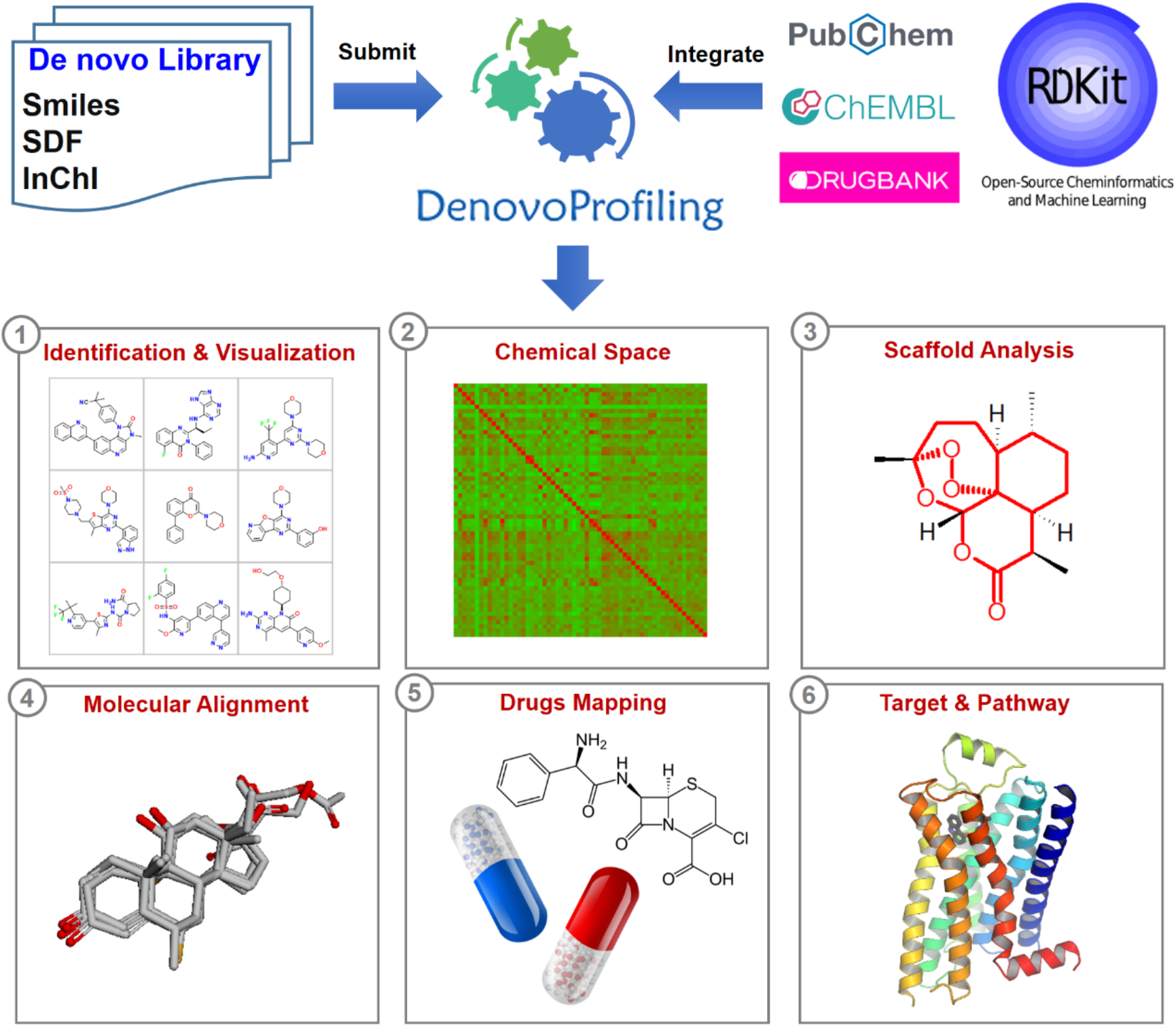
The framework of DenovoProfiling web platform.

### Supported formats

Four widely used chemical formats are supported in DenovoProfiling: SDF, SMILES, InChI, and CDX. All those formats files can be uploaded or the file contents can be pasted and submitted to the web server, except for the binary CDX format, which cannot be pasted. The Open Babel (www.openbabel.org) program was used for chemical file format conversion.

### Modules

Currently, DenovoProfiling provides 6 profiling modules. Each module is functional individually and user could select the module of interest. The implementations for each module are described as follow.

### Identification & Visualization

Identification & Visualization module aims to check whether the de novo structures are already existing and visualize the de novo chemical structures. The submitted de novo molecules were converted into InChIKeys using Open Babel[37]. Subsequently, the InChIKeys were submitted to the PubChem using PubChemPy (https://pubchempy.readthedocs.io), a python package for interacting with PubChem. PubChem is the world’s largest collection of freely accessible chemical information with over 109 million compounds[26]. The PubChem compound ID (cid) were retrieved when de novo molecules were matched. ChemDoodle Web component, a light-weight JavaScript/HTML5 toolkit for chemical graphics, developed by iChemLabs was used for structure visualization[38].

### Chemical Space

Chemical space exploration is an efficient way for de novo library visualization. In this module, the drug-like descriptors were calculated using PaDEL and the distribution were plotted. The chemical similarity heatmap was also generated and was interactive, which user could move or click the cells of the similarity matrix, and the corresponding structures are visualized beside. The principal component analysis was used to visualize the chemical space using PubChem fingerprints.

### Scaffold Analysis

Bemis-Murcko (BM) scaffold[39] was used for scaffold analysis, and implemented by our in-house program[40]. The BM scaffold were generated for each de novo molecule and the number of molecules for each scaffold were calculated and plotted. User could identify the high frequency scaffold conveniently.

### Molecular Alignment

When a de novo molecular library was generated, a straightforward point is to align the focused library to compare the structures of molecules. Molecular alignment module is designed to satisfy this demand. Weighted Gaussian Algorithm (WEGA)[41] was used here for shape alignment. The first molecule was used as the template for alignment. After alignment, user could select the molecules of interest to see or download the alignment results. The three-dimensional conformation alignment was rendered using 3Dmol.js[42].

### Drugs Mapping

Drugs Mapping module was designed to fast retrieve the drugs which are chemical similar to the de novo molecules. The structures of drugs were derived from DrugBank[27]. Inorganic molecules, salts, and duplicates were removed. Similarity between de novo molecule against the drugs were calculated. ECFP fingerprints based Tanimoto similarity coefficient was used.

### Target & Pathway

The bioactivities of the de novo generated molecules were retrieve from ChEMBL database. The target proteins and their bioactivity data were extracted using the generated InChIKeys as query. The retrieved results were summarized in an interactive table and a compound target network. The targeted proteins were further functional enriched using python client of bioinformatics web service DAVID[43]. The enriched KEGG pathways were provided and could be downloaded through an interactive table.

### Web Server Implementation

DenovoProfiling is a publicly accessible platform, which can be accessed through a web browser using the browser server framework. The D3 library of JavaScript (d3js.org/) is used to illustrate the scatterplots, radical plot, and heatmaps. Storage and management of the submitted job data are implemented by MySQL. The back-end server was developed by Golang language. The tools used for constructing the DenovoProfiling are summarized in Supplementary Table S1.

## Results

To test the functionality of DenovoProfiling, we randomly generated a de novo library contains 500 molecules using our previous developed de novo module in DeepScreening[44]. The de novo molecule generation was based on REINVENT[14], an RNN architecture pretrained on more than one million bioactive structures from ChEMBL.

### Structure identification and visualization of de novo library

For a de novo generated library, the first real request is to visualize the chemical structures and know the structure novelty. The Identification & Visualization module was designed to satisfy this demand. The snapshot of this module was shown in Fig. 2. User could browse the structures with mapped PubChem Compound ID (CID). The properties including molecular weight, LogP, HBA, HBD, number of rotatable bonds, TPSA are given by clicking upright plus button. The CID is provided at bottom right and linked the PubChem which provides more detailed compound information.

**Fig. 2.**
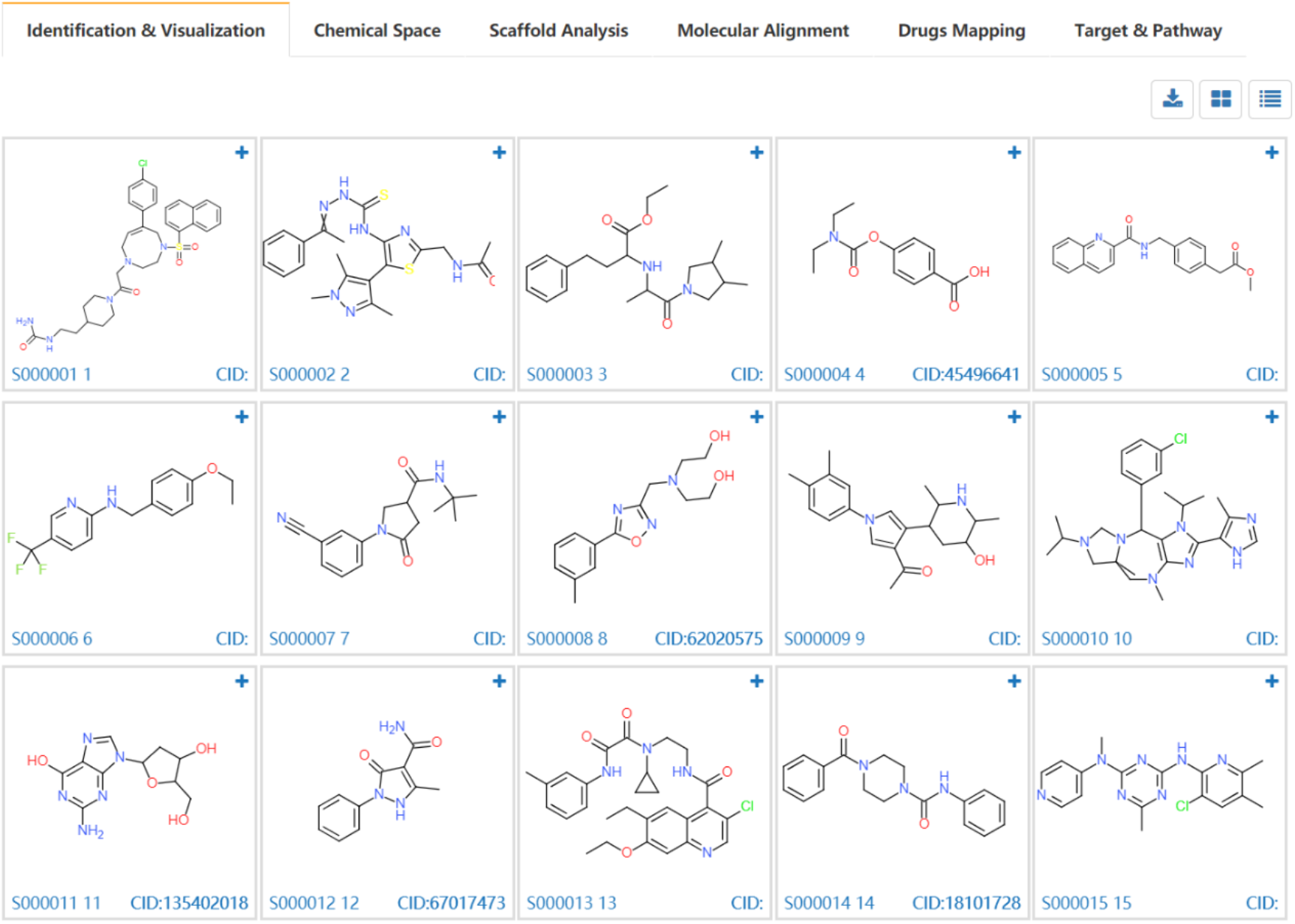
Structure identification and visualization of de novo library.

### Chemical space exploration of de novo library

Chemical structures data are sophisticated, in particular for the de novo generated molecular library, and expert knowledge is highly required[45]. Chemical space visualization is an efficient way to know the structural similarity, or properties similarity of the corresponding molecules through the closeness of the points in this chemical space. Each molecule was defined by a set of numerical descriptors or fingerprints and a set of all molecules was corresponded to the points in the same coordinate-based space. Two important approaches: similarity maps and principal components analysis (PCA) were used in DenovoProfiling. The generated similarity heatmap and PCA plots are interactive, which user could move mouse to the target of point, and the corresponding structure is visualized. The snapshots of similarity map and PCA results were shown in Fig. 3 and Fig. 4. Meanwhile, the distribution of drug-like properties was also plotted, as shown in Fig. 5.

**Fig. 3.**
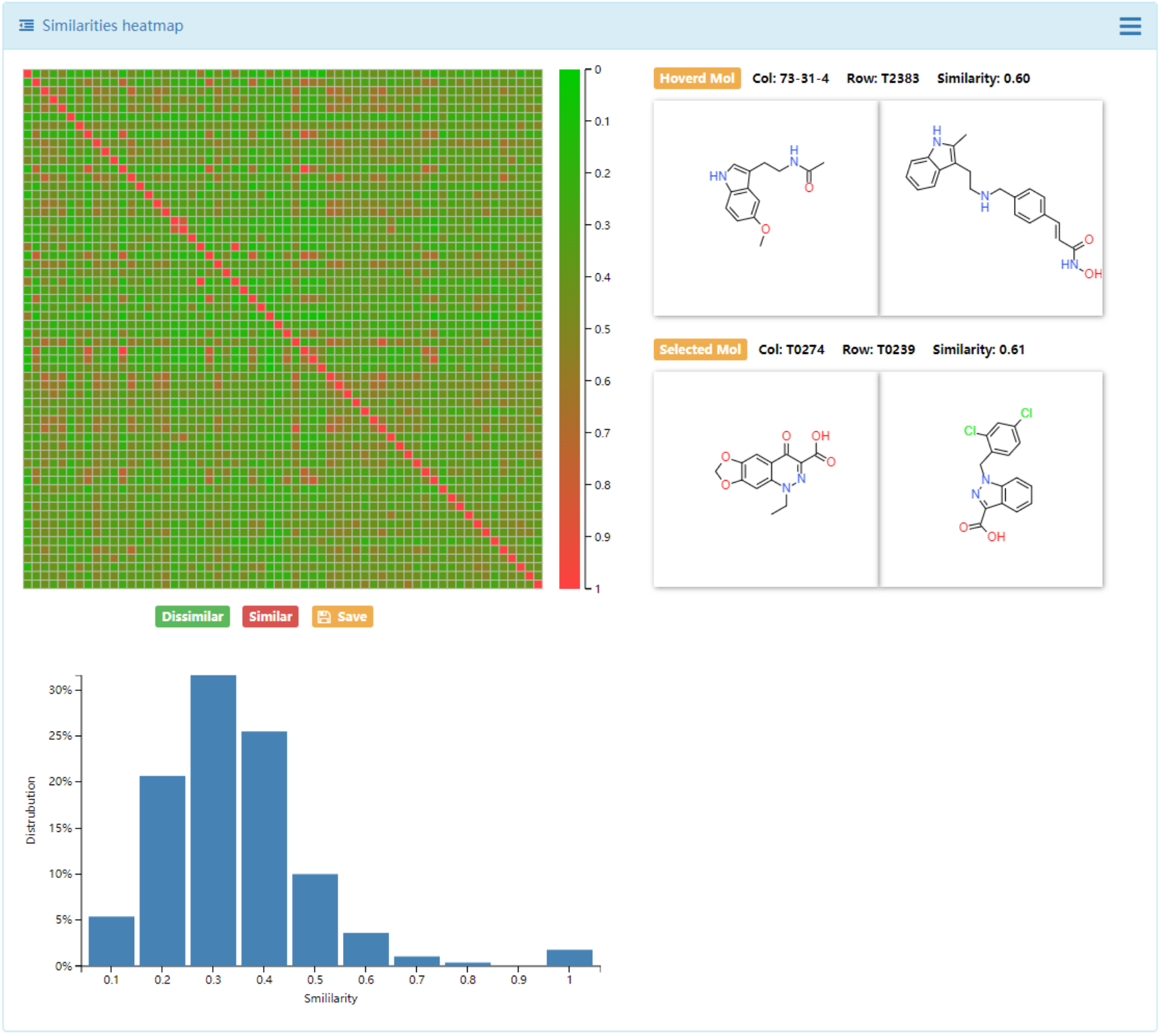
Chemical space illustration using similarity heatmap.

**Fig. 4.**
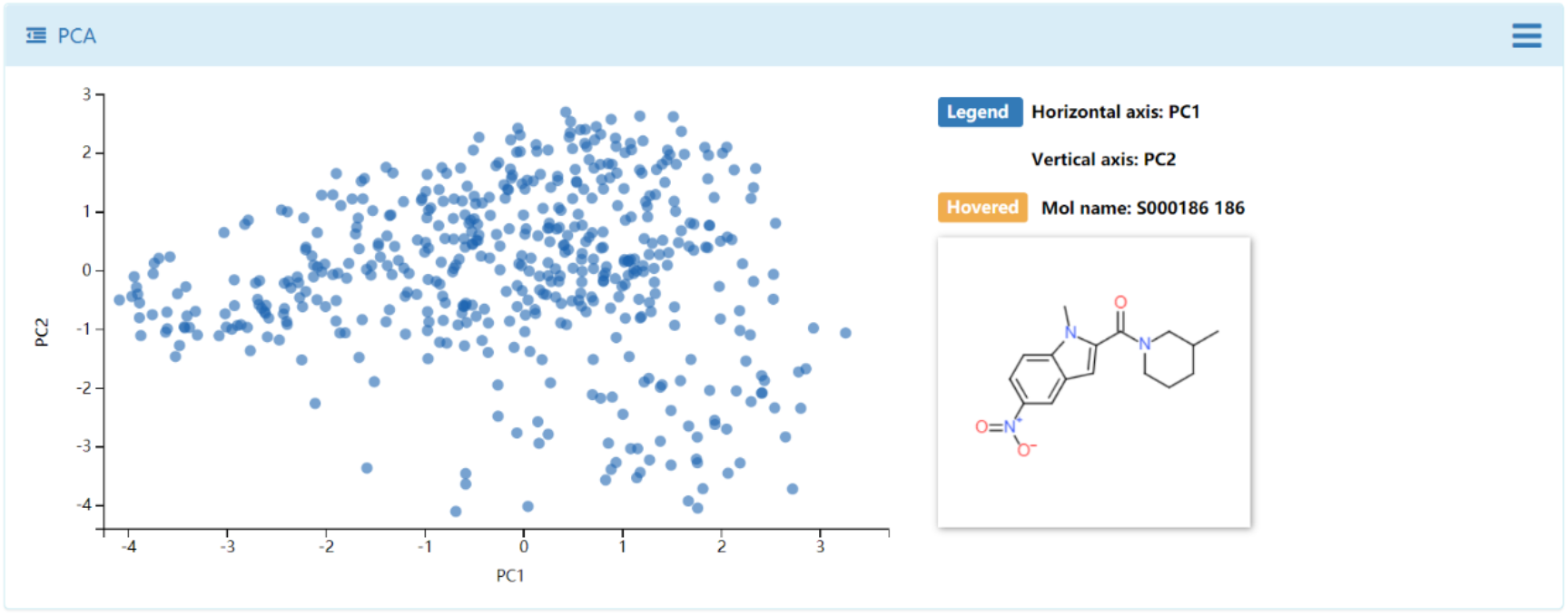
Chemical space illustration using principal component analysis (PCA).

**Fig. 5.**
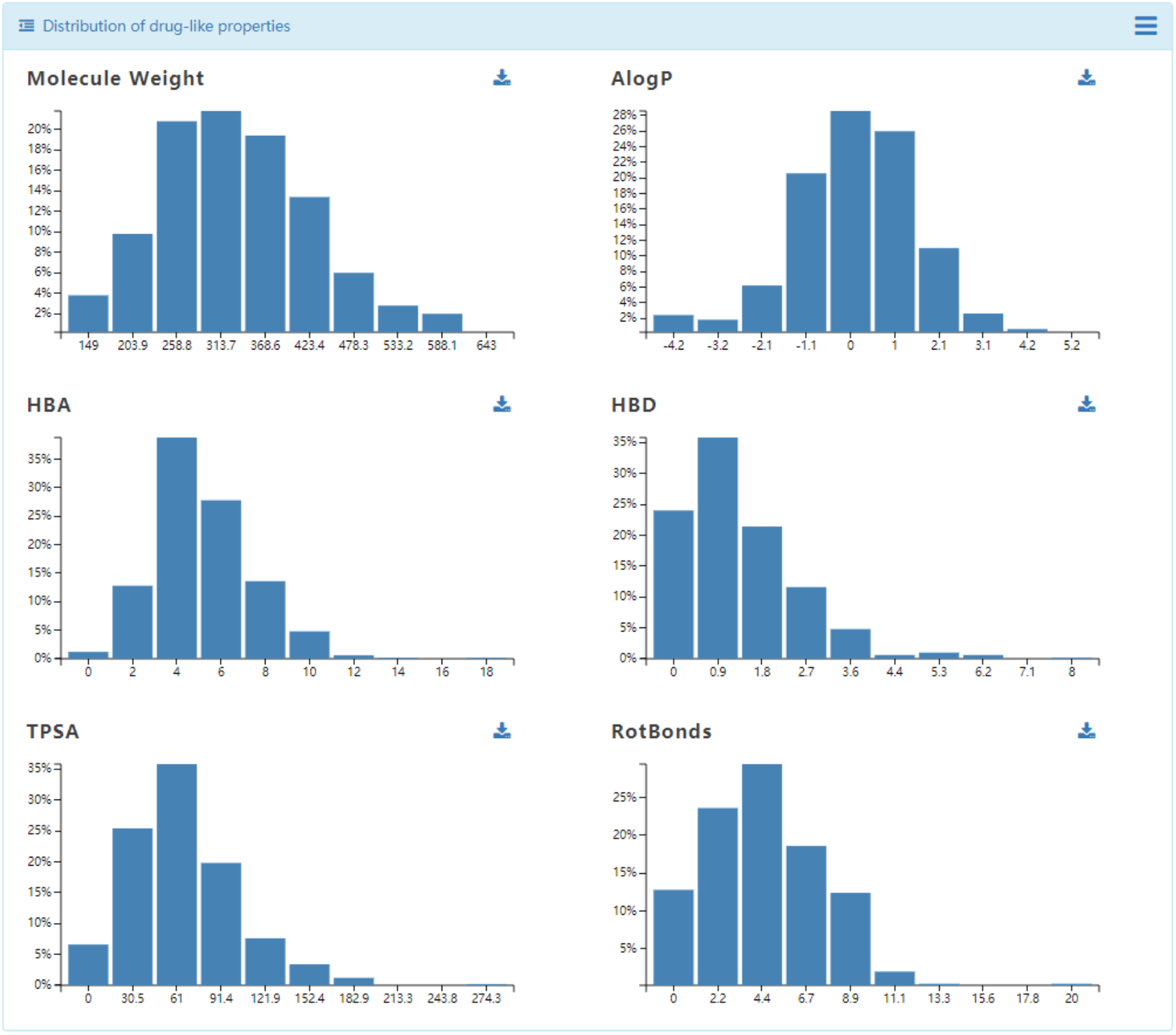
Distribution of drug-like properties.

### Scaffold Analysis of de novo library

Scaffold is an important concept in drug discovery and medical chemistry. Medicinal chemists are seeking chemical with novel scaffolds for a specific biological target[40]. Bemis and Murcko (BM) scaffold [39] approach was used to generate the scaffold for de novo generated library. The complexity and cyclicity of the scaffolds and statistics of each scaffold were interactive illustrated with scatter plot and histogram plot (Fig. 6). As shown in Fig. 7, the structures of scaffolds and their number of molecules were illustrated in grid table. The members of molecules for each scaffold could be browsed by clicking the upright plus button.

**Fig. 6.**
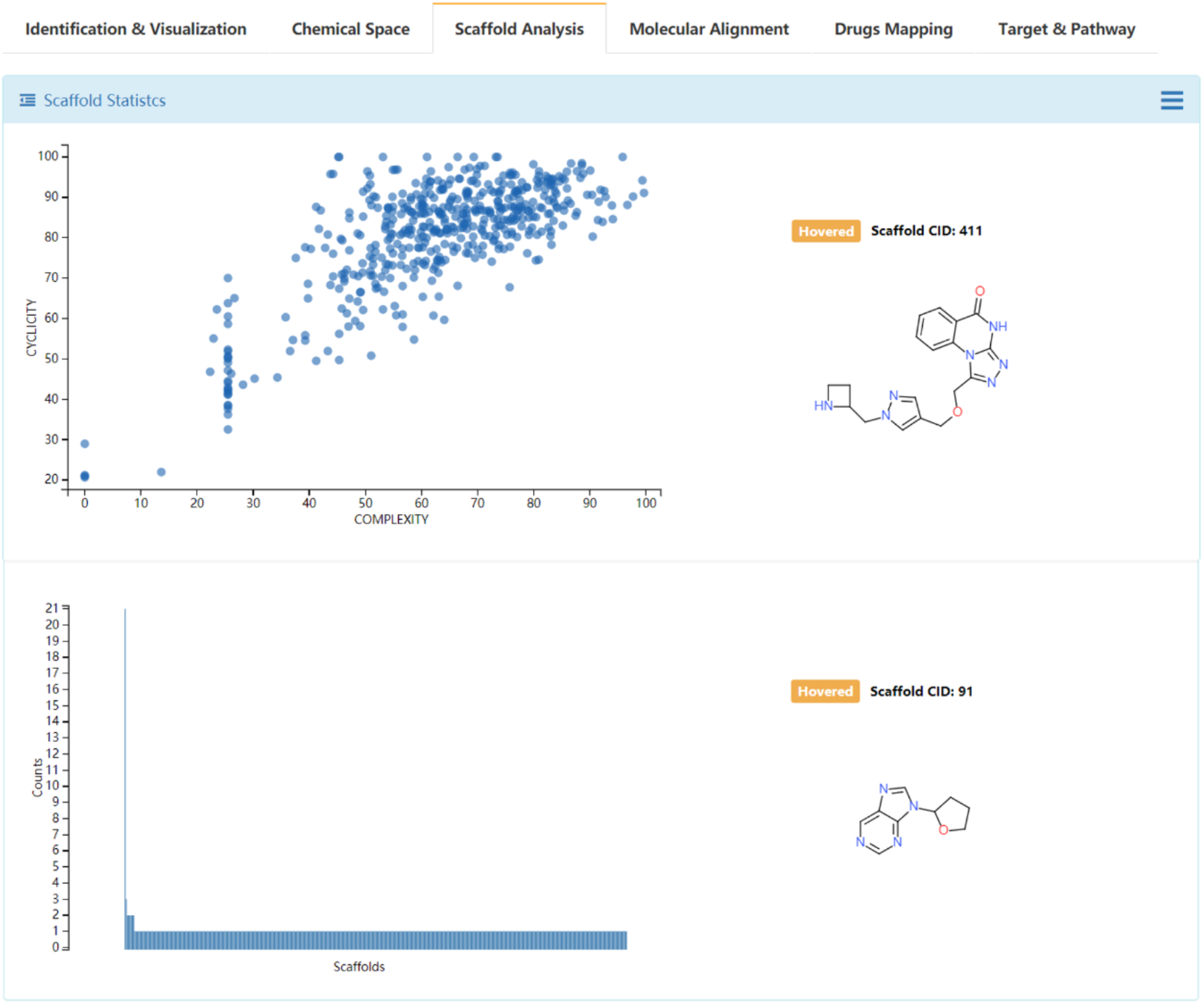
Scaffold statistics of chemical scaffolds of de novo library.

**Fig. 7.**
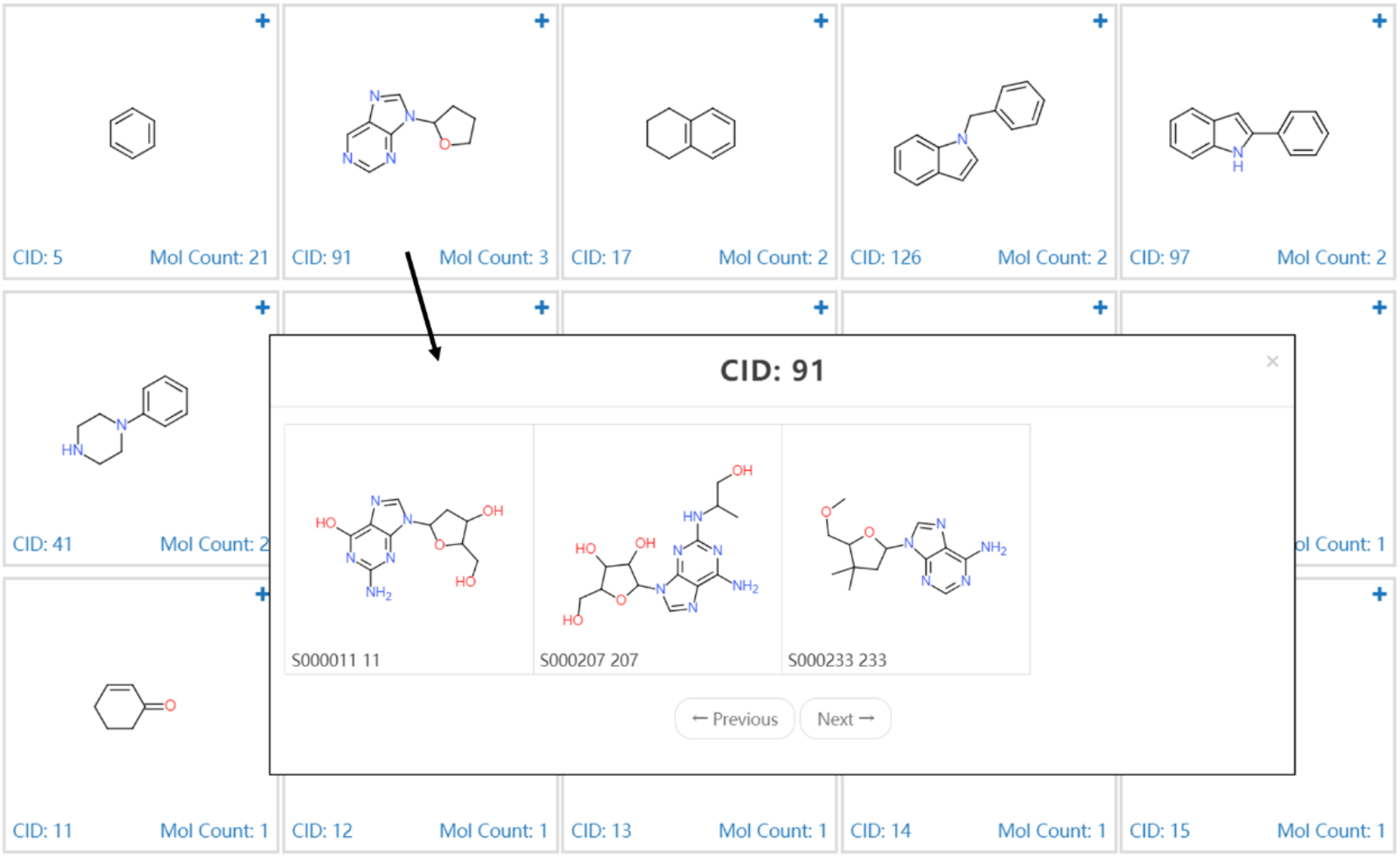
Grid view of the chemical scaffolds of de novo library.

### Molecular alignment of the target-focused or scaffold-constrained library

Generally, medicinal chemist starts from drug target, and attempt to generate a target-focused library, or a scaffold-constraint library for structural optimization. In this case, shape or pharmacophore features based molecular alignment is a good way to compare the difference of the target-focused or scaffold-constraint de novo library. Shape and pharmacophore combined approach in WEGA was used in DenovoProfiling for molecular alignment. User can upload a library of 3D structures, and DenovoProfiling would align all the structures to the first structure of the library. As shown in Fig. 8, user could browse the alignment results, and select the molecules of interest to see the alignment result.

**Fig. 8.**
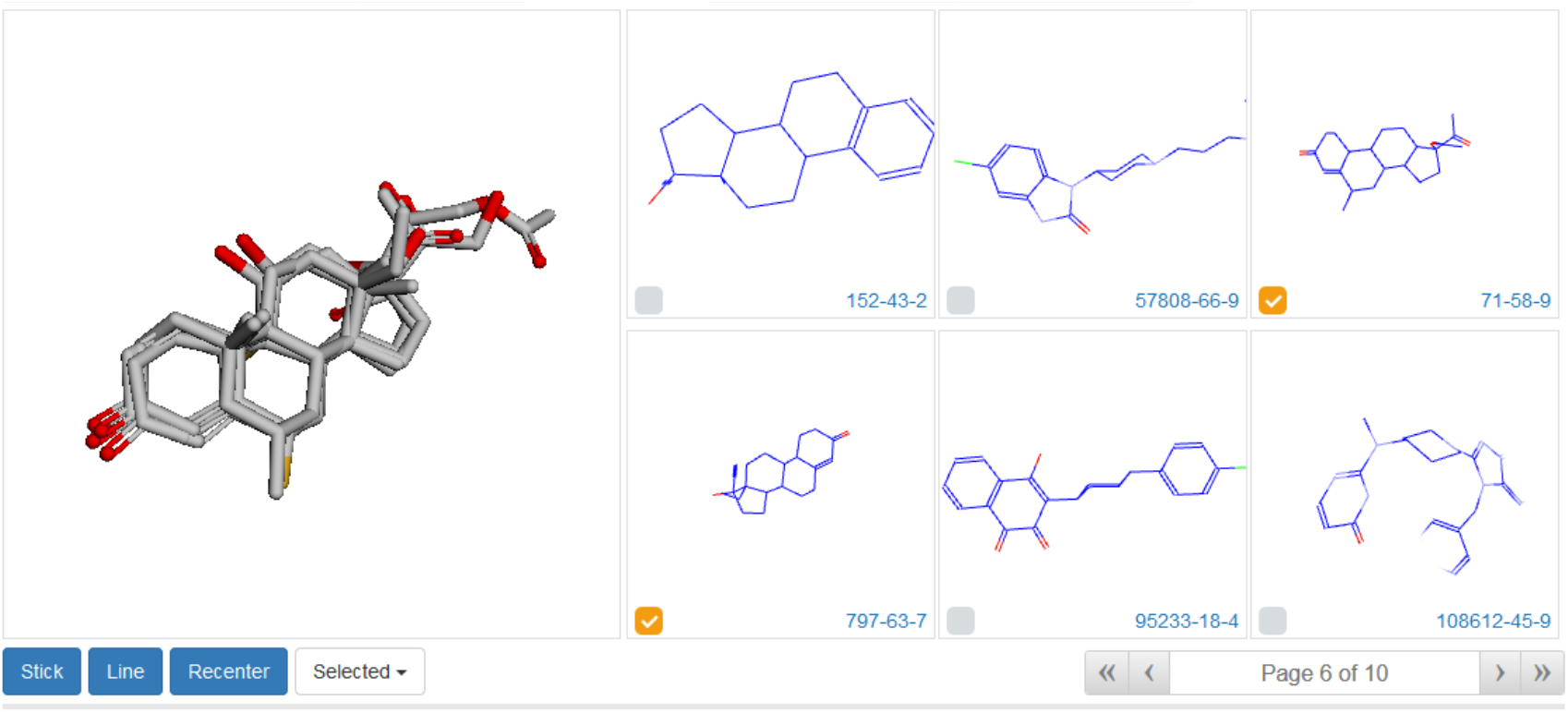
Molecular alignment of the chemical library.

### Drugs mapping of de novo library

De novo generated library usually are randomly and covers a larger chemical space. Though, Identification & Visualization, as mentioned above, identify the structures which have been reported. The structural similar drugs against de novo library are the interest of medicinal chemist. They could fast capture the novelty and pharmacological activities of the de novo compounds when compared with drugs. We prepared an drugs structures library from DrugBank database, and calculate the similarity between the submitted de novo library and drug library. The grid view (Fig. 9) and the table view (Fig. 10) are provided. For the randomly sampled 500 de novo compounds, as shown in Fig. 10, 3 compounds with a maximal drug similarity over 0.9, and their DrugBank ID are also provided and linked to original database. Details of these drugs information could be obtained directly.

**Fig. 9.**
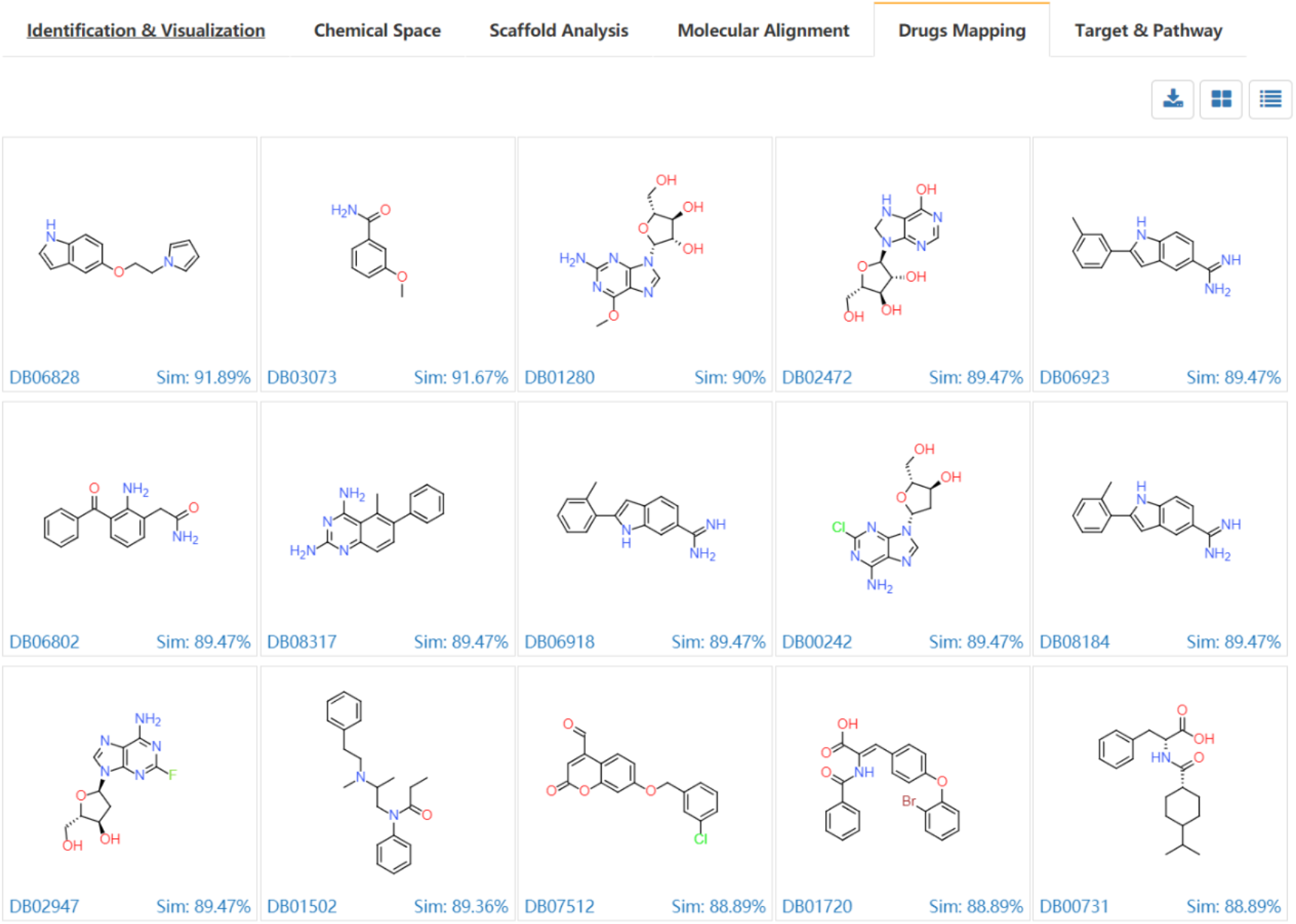
Grid view of the structural similar drugs with de novo molecules.

**Fig. 10.**
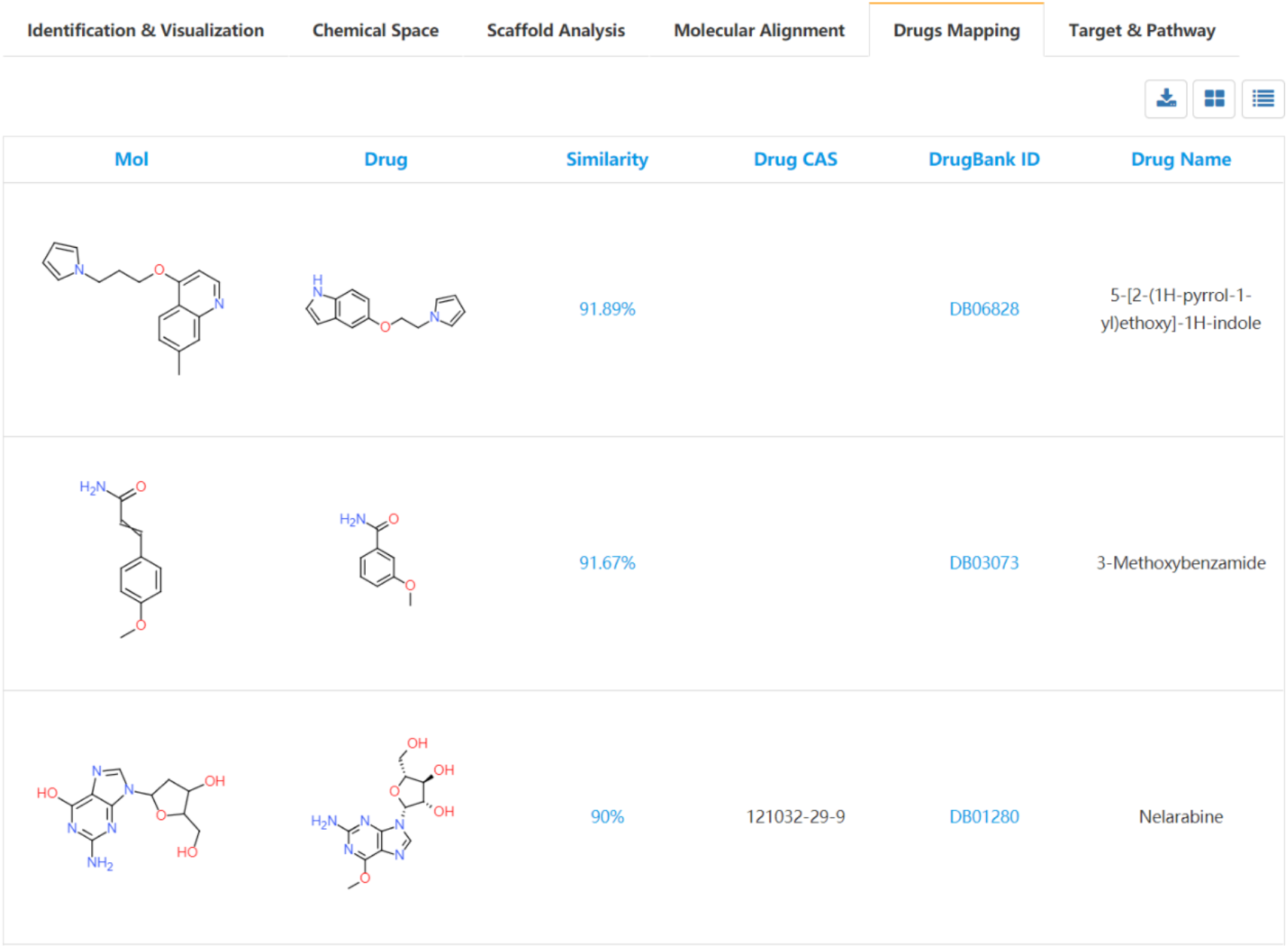
Grid view of the structural similar drugs with de novo molecules.

### Target and pathway profiling for de novo library

The modules we described before are structural annotations for de novo library. Functional profiling is another important part of user concern for de novo library proofing. The ligands and bioactivity data in ChEMBL database were prepared and extracted. The Open Babel was used to generate the unique InChI Key for each structure, then using the InChI Key as query parameter to search against ChEMBL database, the bioactivity data such as Ki, Kd, IC50 and EC50, and corresponding references are extracted. All those results can be analyzed via a user-friendly table view (Fig. 11). Those results are also can be downloaded for local analysis. The compound target relations were further illustrated using compound target network (Fig. 11). The targets are further enriched to pathways and the KEGG pathways are summarized in table (Fig. 12).

**Fig. 11.**
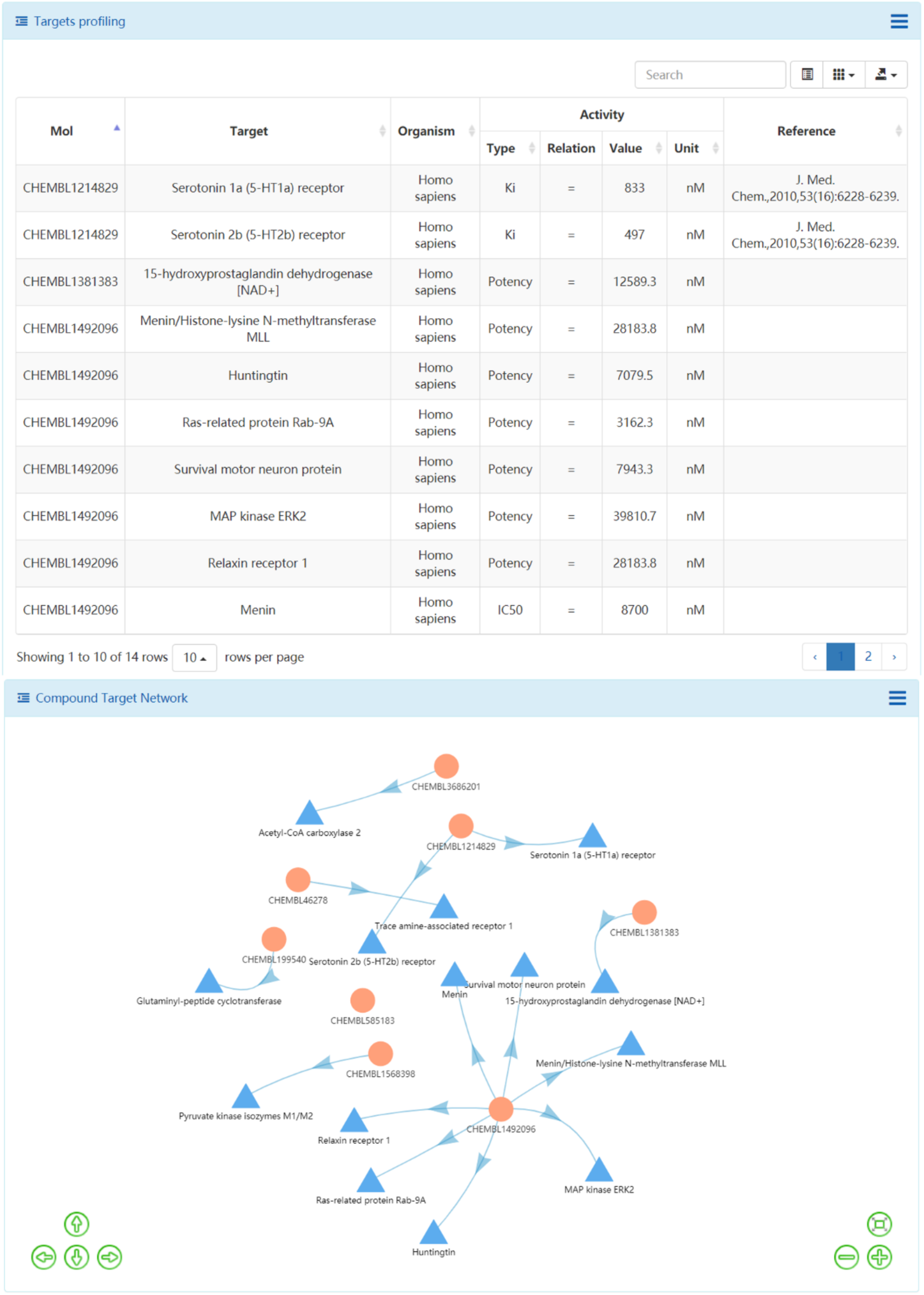
The identified targets in ChEMBL of de novo molecules and the compound target network.

**Fig. 12.**
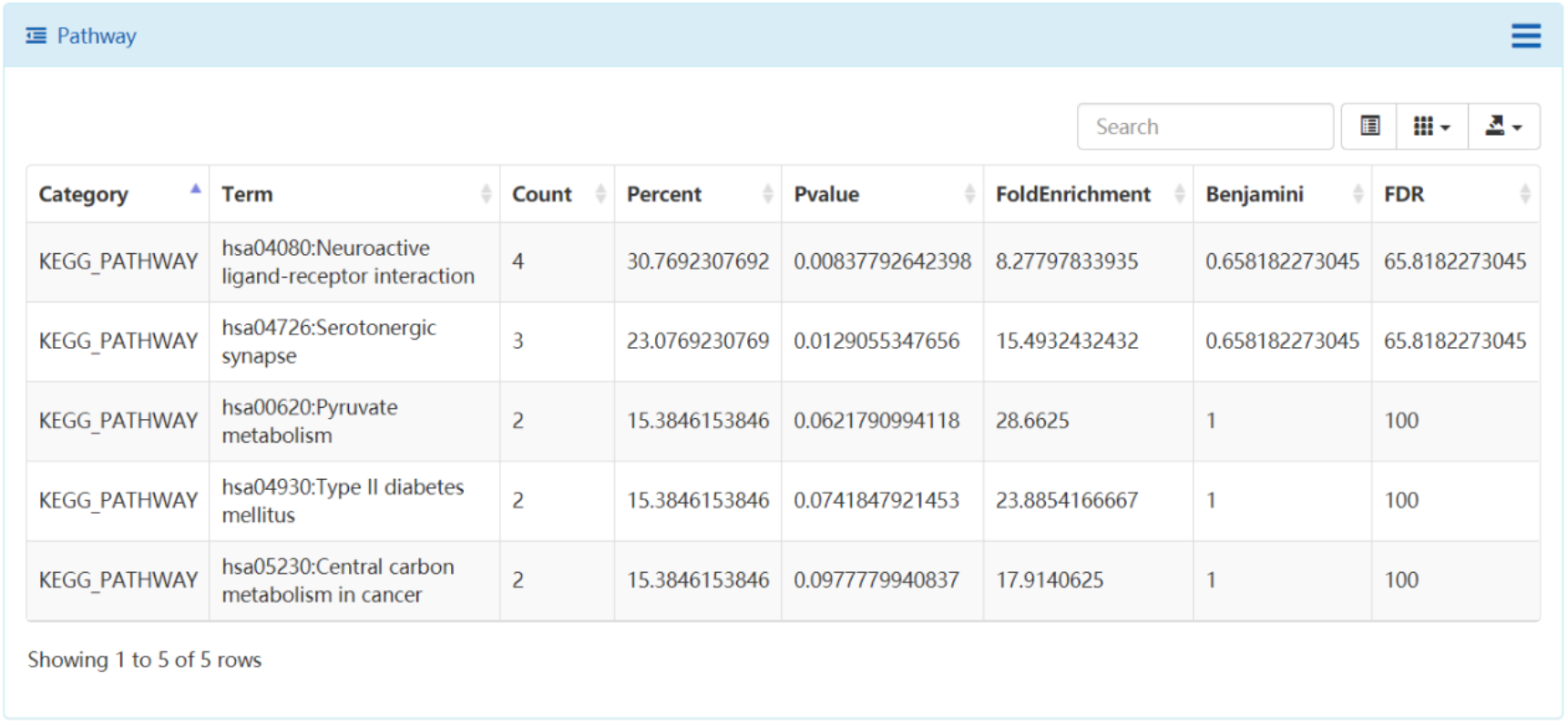
The enriched KEGG pathways using the identified targets in ChEMBL.

## Conclusion

De novo drug design is one of the most promising and scalable approach to accelerate the drug discovery process. Deep learning based de novo molecular generation has shown powerful performance in generating de novo target-focused or property-focused libraries. Fast profiling the de novo generated molecules becomes a practical issue in the de novo drug design. To address this issue, we developed DenovoProfiling, a web-based profiling server for de novo generated molecules. DenovoProfiling support the structure identification, structural visualization, chemical space exploration, scaffold analysis, molecular alignment, drugs profiling, and targets & pathway profiling. These functional modules provide structural and functional annotations for de novo molecules generated from various methods. We believe this web-based tools could facilitate de novo drug design and accelerate the drug discovery.

## Availability and data and materials

The web platform can be accessible at http://denovoprofiling.xielab.net.

## Authors’ contributions

LX and ZL designed the study. ZL and JD implemented the web site. ZL and JD wrote the manuscript. All authors read and approved the final manuscript. We thank professor Jun Xu from Sun Yat-sen University for providing useful suggestions.

## Reference

[1] Segler, M. H. S.; Kogej, T.; Tyrchan, C.; et al. Generating Focused Molecule Libraries for Drug Discovery with Recurrent Neural Networks. ACS Cent. Sci. 2018, 4 (1), 120–131

[2] Mayr, L. M.; Bojanic, D. Novel Trends in High-Throughput Screening. Curr. Opin. Pharmacol. 2009, 9 (5), 580–588

[3] Wildey, M. J.; Haunso, A.; Tudor, M.; et al. High-Throughput Screening. In Annual Reports in Medicinal Chemistry; 2017; pp 149–195

[4] Ripphausen, P.; Nisius, B.; Bajorath, J. State-of-the-Art in Ligand-Based Virtual Screening. Drug Discov. Today 2011, 16 (9–10), 372–376

[5] Zheng, M.; Liu, Z.; Yan, X.; et al. LBVS: An Online Platform for Ligand-Based Virtual Screening Using Publicly Accessible Databases. Mol. Divers. 2014, 18 (4), 829–840

[6] Slater, O.; Kontoyianni, M. The Compromise of Virtual Screening and Its Impact on Drug Discovery. Expert Opin. Drug Discov. 2019, 14 (7), 619–637

[7] Mullard, A. New Drugs Cost US$2.6 Billion to Develop. Nat. Rev. Drug Discov. 2014, 13 (12), 877–877

[8] Yang, X.; Wang, Y.; Byrne, R.; et al. Concepts of Artificial Intelligence for Computer-Assisted Drug Discovery. Chem. Rev. 2019, 119 (18), 10520–10594

[9] LeCun, Y.; Bengio, Y.; Hinton, G. Deep Learning. Nature 2015, 521 (7553), 436–444

[10] Chen, H.; Engkvist, O.; Wang, Y.; et al. The Rise of Deep Learning in Drug Discovery. Drug Discov. Today 2018, 23 (6), 1241–1250

[11] Devi, R. V.; Sathya, S. S.; Coumar, M. S.. Evolutionary Algorithms for de Novo Drug Design – A Survey. Appl. Soft Comput. 2015, 27, 543–552

[12] Schneider, G.; Clark, D. E.. Automated De Novo Drug Design: Are We Nearly There Yet? Angew. Chemie Int. Ed. 2019, 58 (32), 10792–10803

[13] Bian, Y.; Xie, X.-Q. Generative Chemistry: Drug Discovery with Deep Learning Generative Models. 2020, 5276, 1–29

[14] Olivecrona, M.; Blaschke, T.; Engkvist, O.; et al. Molecular De-Novo Design through Deep Reinforcement Learning. J. Cheminform. 2017, 9 (1), 48

[15] Blaschke, T.; Arús-Pous, J.; Chen, H.; et al. REINVENT 2.0: An AI Tool for De Novo Drug Design. J. Chem. Inf. Model. 2020

[16] Blaschke, T.; Olivecrona, M.; Engkvist, O.; et al. Application of Generative Autoencoder in De Novo Molecular Design. Mol. Inform. 2018, 37 (1–2), 1700123

[17] Guimaraes, G. L.; Sanchez-Lengeling, B.; Outeiral, C.; et al. Objective-Reinforced Generative Adversarial Networks (ORGAN) for Sequence Generation Models. arXiv 2017

[18] Langevin, M.; Minoux, H.; Levesque, M.; et al. Scaffold-Constrained Molecular Generation. J. Chem. Inf. Model. 2020, acs.jcim.0c01015

[19] Li, Y.; Hu, J.; Wang, Y.; et al. DeepScaffold: A Comprehensive Tool for Scaffold-Based De Novo Drug Discovery Using Deep Learning. J. Chem. Inf. Model. 2020, 60 (1), 77–91

[20] Yang, Y.; Zheng, S.; Su, S.; et al. SyntaLinker: Automatic Fragment Linking with Deep Conditional Transformer Neural Networks. Chem. Sci. 2020, 11 (31), 8312–8322

[21] Zhavoronkov, A.; Ivanenkov, Y. A.; Aliper, A.; et al. Deep Learning Enables Rapid Identification of Potent DDR1 Kinase Inhibitors. Nat. Biotechnol. 2019, 37 (9), 1038–1040

[22] Yang, Y.; Zhang, R.; Li, Z.; et al. Discovery of Highly Potent, Selective, and Orally Efficacious P300/CBP Histone Acetyltransferases Inhibitors. J. Med. Chem. 2020, 63 (3), 1337–1360

[23] Lipinski, C. A.; Litterman, N. K.; Southan, C.; et al. Parallel Worlds of Public and Commercial Bioactive Chemistry Data. J. Med. Chem. 2015, 58 (5), 2068–2076

[24] Nicola, G.; Liu, T.; Gilson, M. K.. Public Domain Databases for Medicinal Chemistry. J. Med. Chem. 2012, 55 (16), 6987–7002

[25] Burley, S. K.; Berman, H. M.; Bhikadiya, C.; et al. RCSB Protein Data Bank: Biological Macromolecular Structures Enabling Research and Education in Fundamental Biology, Biomedicine, Biotechnology and Energy. Nucleic Acids Res. 2019, 47 (D1), D464–D474

[26] Kim, S.; Chen, J.; Cheng, T.; et al. PubChem 2019 Update: Improved Access to Chemical Data. Nucleic Acids Res. 2019, 47 (D1), D1102–D1109

[27] Law, V.; Knox, C.; Djoumbou, Y.; et al. DrugBank 4.0: Shedding New Light on Drug Metabolism. Nucleic Acids Res. 2014, 42 (D1), D1091–D1097

[28] Mendez, D.; Gaulton, A.; Bento, A. P.; et al. ChEMBL: Towards Direct Deposition of Bioassay Data. Nucleic Acids Res. 2019, 47 (D1), D930–D940

[29] Liu, T.; Lin, Y.; Wen, X.; et al. BindingDB: A Web-Accessible Database of Experimentally Determined Protein-Ligand Binding Affinities. Nucleic Acids Res. 2007, 35 (Database), D198–D201

[30] Bray, S. A.; Lucas, X.; Kumar, A.; et al. The ChemicalToolbox: Reproducible, User-Friendly Cheminformatics Analysis on the Galaxy Platform. J. Cheminform. 2020, 12 (1), 40

[31] Sander, T.; Freyss, J.; Von Korff, M.; et al. DataWarrior: An Open-Source Program for Chemistry Aware Data Visualization and Analysis. J. Chem. Inf. Model. 2015, 55 (2), 460–473

[32] Awale, M.; Probst, D.; Reymond, J. L.. WebMolCS: A Web-Based Interface for Visualizing Molecules in Three-Dimensional Chemical Spaces. J. Chem. Inf. Model. 2017, 57 (4), 643–649

[33] Backman, T. W. H.; Cao, Y.; Girke, T.. ChemMine Tools: An Online Service for Analyzing and Clustering Small Molecules. Nucleic Acids Res. 2011, 39 (SUPPL. 2), 486–491

[34] Deghou, S.; Zeller, G.; Iskar, M.; et al. CART - A Chemical Annotation Retrieval Toolkit. Bioinformatics 2016, 32 (18), 2869–2871

[35] Hilbig, M.; Rarey, M.. MONA 2: A Light Cheminformatics Platform for Interactive Compound Library Processing. J. Chem. Inf. Model. 2015, 55 (10), 2071–2078

[36] Park, S.; Kwon, Y.; Jung, H.; et al. CSgator: An Integrated Web Platform for Compound Set Analysis. J. Cheminform. 2019, 11 (1), 17

[37] O’Boyle, N. M.; Banck, M.; James, C. A.; et al. Open Babel: An Open Chemical Toolbox. J. Cheminform. 2011, 3 (1), 33

[38] Burger, M. C. ChemDoodle Web Components: HTML5 Toolkit for Chemical Graphics, Interfaces, and Informatics. J. Cheminform. 2015, 7 (1), 35

[39] Bemis, G. W.; Murcko, M. A. The Properties of Known Drugs. 1. Molecular Frameworks. J. Med. Chem. 1996, 39 (15), 2887–2893

[40] Liu, Z.; Ding, P.; Yan, X.; et al. ASDB: A Resource for Probing Protein Functions with Small Molecules. Bioinformatics 2016, 32 (11), 1752–1754

[41] Yan, X.; Li, J.; Liu, Z.; et al. Enhancing Molecular Shape Comparison by Weighted Gaussian Functions. J. Chem. Inf. Model. 2013, 53 (8), 1967–1978

[42] Rego, N.; Koes, D.. 3Dmol.Js: Molecular Visualization with WebGL. Bioinformatics 2015, 31 (8), 1322–1324

[43] Huang, D. W.; Sherman, B. T.; Tan, Q.; et al. DAVID Bioinformatics Resources: Expanded Annotation Database and Novel Algorithms to Better Extract Biology from Large Gene Lists. Nucleic Acids Res. 2007, 35 (suppl_2), W169–W175

[44] Liu, Z.; Du, J.; Fang, J.; et al. DeepScreening: A Deep Learning-Based Screening Web Server for Accelerating Drug Discovery. Database 2019, 2019, 1–11

[45] Osolodkin, D. I.; Radchenko, E. V.; Orlov, A. A.; et al. Progress in Visual Representations of Chemical Space. Expert Opin. Drug Discov. 2015, 10 (9), 959–973

